# Contrast-polarity specific mapping optimizes neuronal computation for collision detection

**DOI:** 10.1101/2022.05.10.491323

**Authors:** RB Dewell, Y Zhu, M Eisenbrandt, R Morse, F Gabbiani

## Abstract

Neurons receive information through their synaptic inputs, but the functional significance of how those inputs are mapped on to a cell’s dendrites remains unclear. We studied this question in a grasshopper visual neuron that tracks approaching objects and triggers escape behavior before an impending collision. In response to black approaching objects, the neuron receives OFF excitatory inputs that form a retinotopic map of the visual field onto compartmentalized, distal dendrites. Subsequent processing of these OFF inputs by active membrane conductances allows the neuron to discriminate the spatial coherence of such stimuli. In contrast, we show that ON excitatory synaptic inputs activated by white approaching objects map in a random manner onto a more proximal dendritic field of the same neuron. This random synaptic arrangement results in the neuron’s inability to discriminate the coherence of white approaching stimuli. Yet, the neuron retains the ability to discriminate stimulus coherence for checkered stimuli of mixed ON/OFF polarity. The coarser mapping and processing of ON stimuli thus has a minimal impact, while reducing the total energetic cost of the circuit. Further, we show that these differences in ON/OFF neuronal processing are behaviorally relevant, being tightly correlated with the animal’s escape behavior to light and dark stimuli of variable coherence. Our results show that the synaptic mapping of excitatory inputs affects the fine stimulus discrimination ability of single neurons and document the resulting functional impact on behavior.

## Introduction

A major goal of neuroscience is determining the mechanisms of neural computation – how is information processed within neural circuits and how do cellular properties produce these abilities. Neurons receive information about the external world through the pattern of their synaptic inputs. So, determining how they integrate them is critical. For some tasks, such as assessing the presence of an impending threat, sensory discriminations must be quick and reliable for survival. In neural processing there are trade-offs, though, between speed, high-resolution discrimination, and energy efficiency (Attwell and Laughlin, 2001; Laughlin, 2001; Vincent and Baddeley, 2003; Hasenstaub et al., 2010). Evolution presumably settles on solutions that minimize energy expenditure while maintaining required processing abilities.

Individual neurons process information by discriminating between patterns of synaptic inputs impinging on their dendrites. Pre-synaptic targeting, dendritic morphology, compartmentalization and active membrane properties all shape neuronal processing (London and Hausser, 2005; Major et al., 2013; Lefebvre et al., 2015; Hawkins and Ahmad, 2016). Many neurons, including the principal excitatory neurons of the mammalian cortex, pyramidal neurons, have distinct dendritic subfields proximal and distal to the site of spike initiation that receive functionally segregated synaptic inputs processed by distinct membrane conductances (Behabadi et al., 2012; Major et al., 2013; Hawkins and Ahmad, 2016). In these neurons, the role of anatomical segregation within dendrites in integrating inputs from distinct neural pathways remains largely unclear.

In the visual system, neural channels transmitting information about luminance increases (ON) and decreases (OFF) constitute one key example of parallel pathways requiring integration (Chen et al., 2017; Williams et al., 2021). Although luminance information is segregated between ON and OFF cells early on in the retina of insects and mammals, real world scenes involve a mix of both (Leonhardt et al., 2016; Chen et al., 2019; Mazade et al., 2019; Mulholland and Smith, 2021). The visual world contains more information in luminance decreases, particularly for smaller, faster objects (Simoncelli and Olshausen, 2001; Clark et al., 2014; Leonhardt et al., 2016; Chen et al., 2019). Although, the contrast statistics of approaching predators has not been determined, it is likely that they also contain more OFF than ON information (Zhou et al., 2022). How these pathways are integrated within individual neurons driving behavior remains unanswered, though.

To address these questions, we leveraged a well-studied, identified neuron in grasshoppers that receives ON and OFF inputs across three distinct dendritic subfields. It processes these inputs to detect impending collision, resulting in a spiking output critical to the generation of escape behavior (O’Shea and Williams, 1974; Gabbiani et al., 1999; Fotowat et al., 2011; Dewell and Gabbiani, 2018a). This so-called lobula giant movement detector (LGMD) neuron receives excitation and inhibition from all ommatidia (facets) of the compound eye after processing in two intermediate neuropils. The LGMD is about as large as a human cerebellar Purkinje cell and its output is faithfully relayed by a second neuron, the descending contralateral movement detector (DCMD), to the cells coordinating escape jumps. The LGMD distinguishes the coherence of dark (OFF) approaching objects through a retinotopic synaptic input mapping onto compartmentalized dendrites, and through intracellular processing by differentially distributed active conductances (Dewell and Gabbiani, 2018a; Zhu et al., 2018). These include an H-conductance and an inactivating K^+^ conductance localized in dendrites, as well as calcium and calcium sensitive K^+^ conductances localized close to the spike initiation zone, and a M-type K^+^ conductance presumably localized in the axon (Peron and Gabbiani, 2009; Dewell and Gabbiani, 2018a, 2018b). Little is known on the LGMD processing of ON inputs.

Using calcium imaging, electrophysiology, pharmacology, and modeling, we characterized a novel excitatory pathway to the LGMD for ON inputs. We show that the mapping of ON and OFF excitation to distinct dendritic fields has behavioral consequences. Further, we show that the mechanisms of ON-OFF integration within the LGMD’s dendritic arbor allows the detection of approaching objects of any contrast polarity, while maintaining in most cases selectivity for their spatial coherence. Finally, we show that the mapping of ON and OFF excitatory inputs on distinct dendritic subfields allows the implementation of coherence discrimination in an energy efficient manner, suggesting integration principles likely applicable to other neurons, including within our own brains.

## Results

Visual circuits are subdivided into ON and OFF pathways, but animals can face approaching threats that are light, dark, or a mottled combination. Although extensive work has investigated the role of OFF pathways in detecting impending collision, how ON and OFF signals are integrated to decide whether to initiate escape is unknown. We used grasshoppers which have well characterized jump escape behaviors initiated by an identified neuron in the optic lobe to explore the neuronal integration of ON and OFF contrast polarities for collision avoidance.

### Jump probability, but not timing, is independent of contrast polarity

In grasshoppers and other species there has also been extensive research on the behavioral responses produced by black looming stimuli, but there has been much less investigation into how behavior depends on contrast polarity although mice show a clear behavioral preference for simulated black approaching stimuli (Yilmaz and Meister, 2013). First, we tested whether the animals escape from simulated white objects approaching at constant speeds as they do for black ones (Fig. 1A; Video 1). These looming stimuli are characterized by the parameter *l*/|*v*| (i.e., the ratio of their half-size to approach speed, see Fig. 1A; Gabbiani et al., 1999). Grasshoppers jumped similarly in response to all looming stimuli, with escape jumps from 57% of black (OFF) looms and 45% of white (ON) looms (Fig. 1B). For neither contrast polarity did the approach speed influence jump probability (for a fixed stimulus size; p = 0.77 for white, p = 0.42 for black, KW). The jump timing changed slightly with contrast polarity, however, with white stimuli producing earlier jumps for intermediate approach speeds. (Fig. 1C). For looming stimuli with a *l*/|*v*| of 80 ms the median jump time was 223 ms earlier (p = 8.9 • 10^−4^); the jump time was not significantly earlier for white looms with a *l*/|*v*| of 40 or 120 ms.

**Figure 1.**
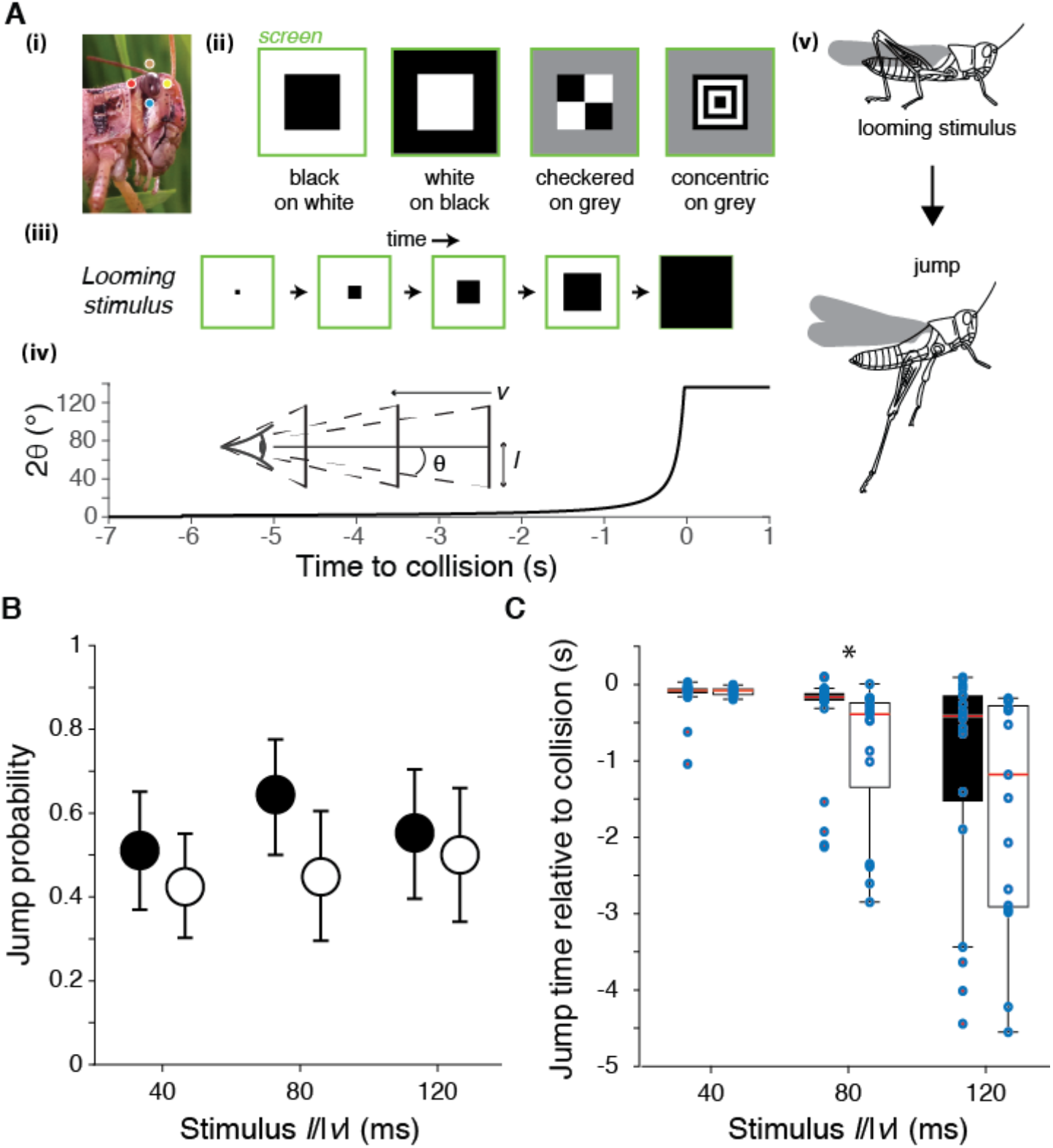
Escape behavior to white and black looming stimuli. A) The locust eye (i) was stimulated using four simulated approaching squares (looming stimuli) with different contrast polarities (ii). (iii), (iv): Schematic of looming stimulus expansion on the retina (in the case of a black square) and time course of angular expansion for *l*/|*v*| = 80 ms, respectively. Inset illustrates the definition of the approach speed (*v* < 0), the square half-size (*l*) and the half-angle of the stimulus subtended at the retina (θ). Behaviorally, such stimuli lead to escape jumping (v). In (i), colored dots match those of Fig. 2A. B) Grasshoppers consistently jumped to white and black stimuli of different *l*/|*v*|. White stimuli had slightly lower, but not significantly reduced escape responses (p = 0.065, Fisher’s test). Error bars are 95% confidence intervals. C) White stimuli produced jumps earlier relative to collision than did black stimuli for *l*/|*v*| of 80 ms; for *l*/|*v*| of 40, 80, and 120 ms, p = 0.83, 8.9 • 10^−4^, and 0.26 respectively (WRS). Displayed as median ± mad with points showing individual trials. For B and C, N = 263 trials from 7 grasshoppers. WRS: Wilcoxon rank-sum test.

### Dendritic field segregation of ON/OFF excitatory inputs to the LGMD neuron

Previous investigations into the visual detection of impending collisions have focused on black looming stimuli, generating a detailed characterization of their processing by the presynaptic circuitry to the LGMD and by active membrane conductances within the LGMD. The LGMD has a large distal dendritic field (field A) and two smaller ones more proximal to the spike initiation zone (SIZ; fields B and C; Fig. 2A). Inhibitory inputs to the LGMD arise from pathways segregated by contrast polarity, with OFF inhibition impinging on dendritic field C and ON inhibition impinging on field B (Rowell et al., 1977). The LGMD’s excitatory inputs, however, were believed to be processed independent of polarity, with both ON and OFF pathways projecting retinotopically to field A (Fig. 1A (i), Fig. 2A, colored dots; O’Shea and Rowell, 1976). This arrangement has been documented for OFF, but not for ON stimuli (Peron et al., 2009; Zhu and Gabbiani, 2016). We tested its validity for ON inputs with wide-field calcium imaging of all three dendritic fields during presentation of light and dark looming stimuli (Fig. 2).

**Figure 2.**
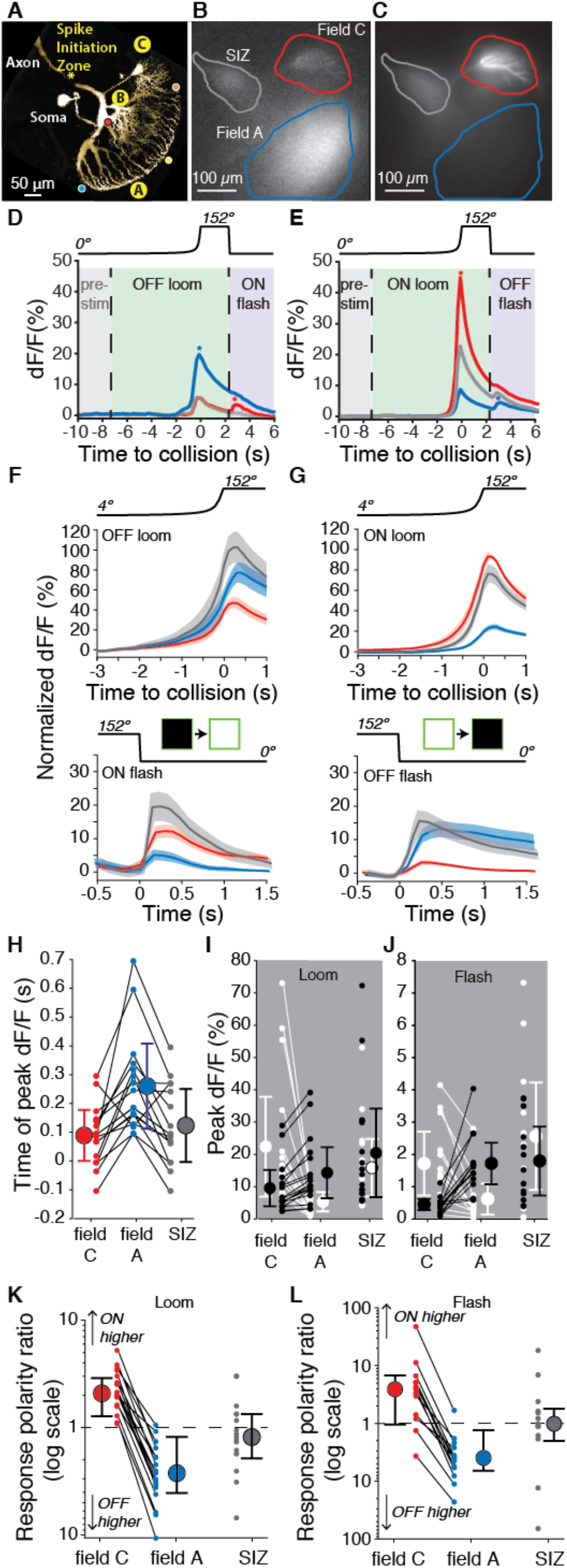
Luminance increases produce excitation in dendritic field C of the LGMD, unlike field A. A) Micrograph (2-photon scan) of the LGMD neuron illustrating the neurites imaged: dendritic fields A and C, as well as the spike initiation zone (SIZ). Colored dots matching those of Fig. 1A (i) indicate retinotopic mapping of visual field onto LGMD dendritic field A. B, C) Micrographs of maximum dF/F taken during presentation of OFF and ON, looming stimuli. Micrographs rotated clockwise ∼120º relative to A. Areas enclosed by solid lines are ROIs used to compute dF/F time course. D, E) Corresponding dF/F time course in response to an OFF, respectively ON, looming stimulus within the ROIs marked by matching colored lines in B, C. Abbreviations: pre-stim, baseline prior to looming; ON flash, disappearance of full-size black square from screen. Field A showed a larger increase in dF/F during looming (blue star, D). Field C showed a larger rebound to the ON flash (red star, D). Conversely, the ON loom produced a larger dF/F in field C (red star, E). Its disappearance (OFF flash) produced a larger rebound dF/F in field A (blue star, E). F) Population average normalized dF/F for the 3 ROIs during presentation of black stimuli (same color code as in B). Lines and shaded regions are mean ± s.e.m. In each animal, normalization is to the peak response for ON looms. G) Similar average normalized dF/F for ON stimuli. H) The peak dF/F in field A occurred later than in field C for both ON and OFF looms (p = 2.7 • 10^−4^). I) Peak dF/F response to ON and OFF looming stimuli for each animal. Responses to ON stimuli were higher in field C (p = 1.5 • 10^−5^) and black looming responses were higher in field A (p = 1.5 • 10^−5^). J) Peak dF/F response to post-loom flash. Responses to ON flashes were higher in field C (p = 2.4 • 10^−4^) while OFF flash responses were higher in field A (p = 2.4 • 10^−4^). K) For all animals, loom response polarity is biased towards ON stimuli in field C (p = 1.5 • 10^−5^). In field A, it is biased towards OFF stimuli in 16 of 17 animals (p = 2.7 • 10^−4^). L) Similarly, response polarity is biased towards ON flashes in field C (p = 2.3 • 10^−3^), and towards OFF flashes in field A (p = 2.7 • 10^−4^). For F-L N = 17, and colors in D-H and K, L match the ROIs shown in (B) – SIZ is grey, field A is blue, and field C is red. The p-values for H-L are from two-sided sign tests.

The previously described excitatory synaptic inputs onto field A are mediated through nicotinic acetylcholine receptor (nAChR) channels that are calcium permeable (Peron et al., 2009). As the dendrites lack voltage-gated Ca^2+^ channels, this allows direct imaging of synaptic activation with an intracellularly injected fluorescent calcium dye, while voltage-gated Ca^2+^ channels near the SIZ allow imaging of the neuron’s SIZ activation (Peron and Gabbiani, 2009; Peron et al., 2009). Presentation of an OFF-looming stimulus produced the expected increase in fluorescence in field A (Fig. 2B). In response to an ON-looming stimulus, however, fluorescence increased in field C and not in field A (Fig. 2C; Video 2). No fluorescence changes were observed in field B, regardless of stimulus contrast polarity. The time course of the fluorescence increase was similar in fields A and C and near the SIZ with a clear difference in dF/F amplitude depending on contrast polarity (Fig. 2D, E; Video 3,4). This provides the first evidence that excitation from ON and OFF inputs are segregated and independently integrated within the LGMD.

For comparison between animals, we normalized each animal’s fluorescence signal to the maximum response to an ON loom in field C. During OFF-looming expansion, field A consistently had a larger response, while field C responded more to ON looms (Fig. 2F). Following looming stimuli, the screen changed abruptly back to its original background luminance after ∼2 s causing a ‘flash’ of the opposite polarity to the loom. ON flashes produced a larger response in field C and OFF flashes a larger response in field A (Fig. 2D, F). For looms of either polarity, the peak fluorescence change occurred near the projected time of collision, with the peak response in field A trailing the peak fluorescence in field C and the SIZ (Fig. 2H).

To characterize the consistency in polarity segregation across animals, we compared peak responses between dendritic fields for ON and OFF stimuli (Fig. 2I, J) and the ratio of peak responses between ON and OFF stimuli within each dendritic field (Fig. 2K, L). ON looms produced a 2.4 ± 1.2 times larger response in field C than did OFF looms, while in field A OFF loom responses were 3.6 ± 1.1 times those of ON looms (mean ± s.d.). Flash responses showed a similar preference with responses 3.4 ± 1.7 times higher for ON than for OFF flashes in field C, and responses 5.6 ± 2.2 times higher for OFF than ON flashes in field A. Calcium fluorescence at the SIZ did not consistently differ with stimulus polarity for either loom or flash stimuli (Fig. 2I-L).

### ON field C synaptic inputs are mediated by nicotinic acetylcholine receptors

Having discovered an unexpected ON pathway projection onto field C of the LGMD, we tested how similar the ON inputs to field C are to the previously characterized OFF inputs in field A. Since the field A inputs are mediated by nAChRs, we examined whether that was also the case for the ON inputs to field C. Direct iontophoresis of acetylcholine (ACh) to field C dendrites increased calcium fluorescence in parallel with the amount of applied ACh (Fig. 3A, B). That the field C receptors are nicotinic was confirmed with the use of the nAChR antagonist mecamylamine. Local puffing of mecamylamine near field C reduced calcium fluorescence produced by looming stimuli within field C and at the SIZ (Fig. 3C-F). Mecamylamine reduced peak dF/F in field C by 80 ± 8.5% for ON looms and 75 ± 17% for OFF looms (mean ± s.d.).

**Figure 3.**
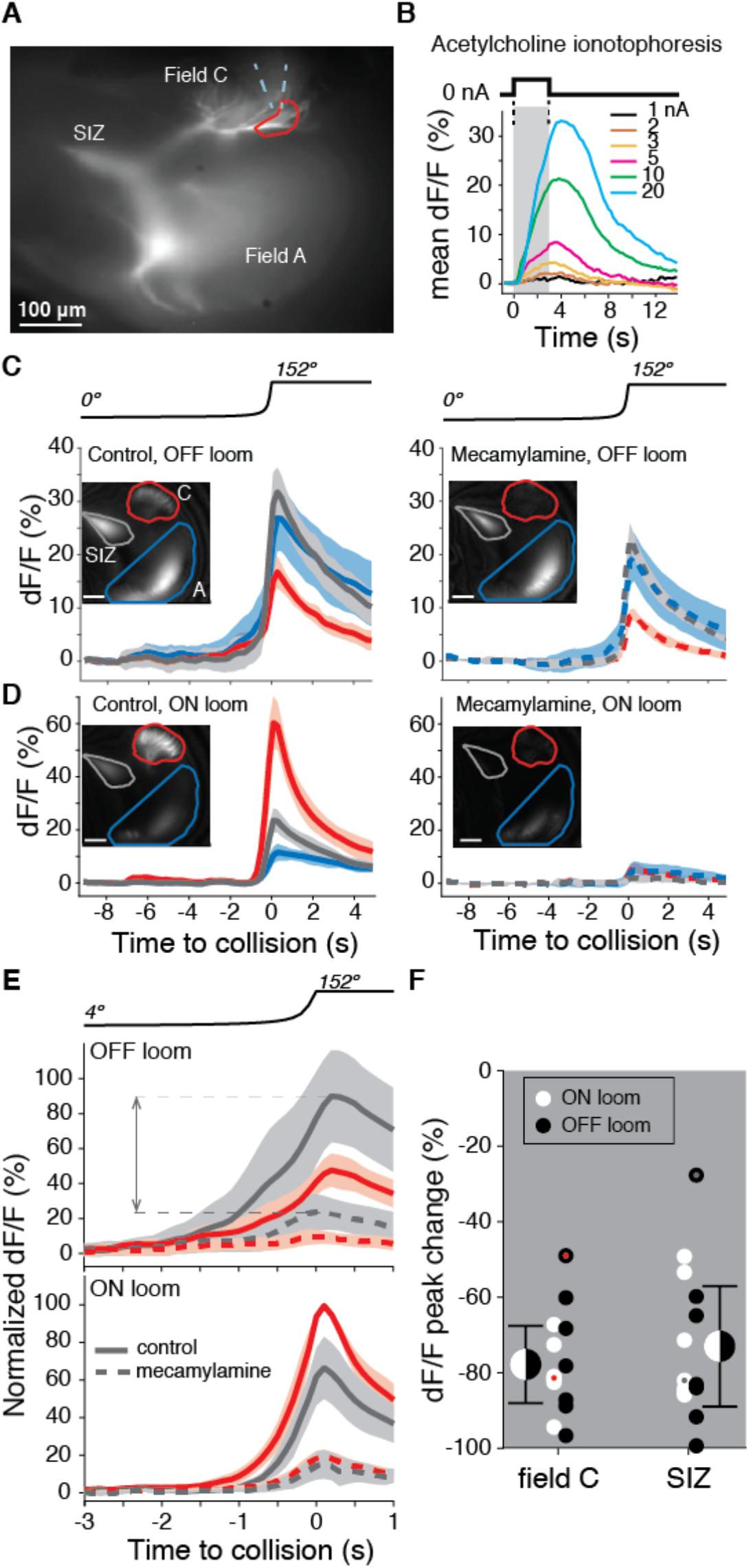
Dendritic field C receives excitatory synaptic inputs through nAChRs. A) Micrograph of the LGMD stained with OGB1 showing the locations of the spike initiation zone (SIZ), dendritic field A (both out of focus), and dendritic field C (in focus). Dashed blue wedge: location of iontophoresis electrode; closed red curve: boundary of area used to compute dF/F in B) Example dF/F to ACh iontophoresis for square current pulses (1 – 20 nA; top inset and grey shading). C, D) Left: example responses from one animal showing the dF/F produced by Ca^2+^ influx in response to an OFF (C) or ON (D) looming stimulus in all three LGMD subregions (insets, as in A). Right: application of the nAChR blocker mecamylamine to field C reduced this influx (pipettes were placed as shown in A). Lines and shaded region are trial mean ± s.d. E) Average dF/F (± s.e.m.) of 7 animals before (solid lines) and after mecamylamine application (dashed lines) in field C (red) and at the SIZ (grey). Responses were normalized to each animal’s mean peak dF/F response in field C during a ON looming stimulus. Dashed horizontal lines and double arrow show how the peak change plotted in F was computed. F) Looming responses were reduced in field C by 78% (p = 0.015, ST) and by 73% at the SIZ (p = 0.015, ST) on average. Field A dF/F was reduced as well, 58%, but less than field C (p = 0.013, ST). Black and white dots: individual animal responses. Half black and white discs: mean ± s.d. pooled across animals and stimuli. Dots marked red (field C) and grey (SIZ) correspond to the animal shown in C, D. ST: sign test.

### ON field C synaptic inputs are randomly distributed

Excitatory inputs impinging on field A follow a precise retinotopic arrangement (Peron et al., 2009; Zhu et al., 2018). To test whether this was also the case for field C excitation, we examined the timing of activation within dendritic subregions (Fig. 4A). Both ON and OFF looming stimuli produced synchronous activation of field C dendrites (Fig. 4B, C). This differs from field A which exhibits sequential activation of dendritic regions as the loom expands (Fig. 4D; Zhu et al., 2018). This difference between the two dendritic fields was even more pronounced for the responses to translating squares. Field A showed sequential activation of subregions as the visual object translated across the LGMD receptive field (Fig. 4E; Video 7). Within field C, however, all dendritic subregions were activated simultaneously (Fig. 4F; Video 5 and 6). This was confirmed by looking at the range of peak times across subregions (Fig. 4G). Despite field C being subdivided into more dendritic subregions than field A, thus biasing it toward a larger range, it consistently had more synchronous activation of dendritic regions than field A. For 12 of 18 animals, all field C subregions had the same dF/F peak time in response to ON looms, and in no experiment did the responses suggest a retinotopic mapping of inputs to field C. Thus, the synaptic mapping of ON inputs onto field C is strikingly different from the previously described, ordered retinotopic organization of OFF synaptic inputs onto field A.

**Figure 4.**
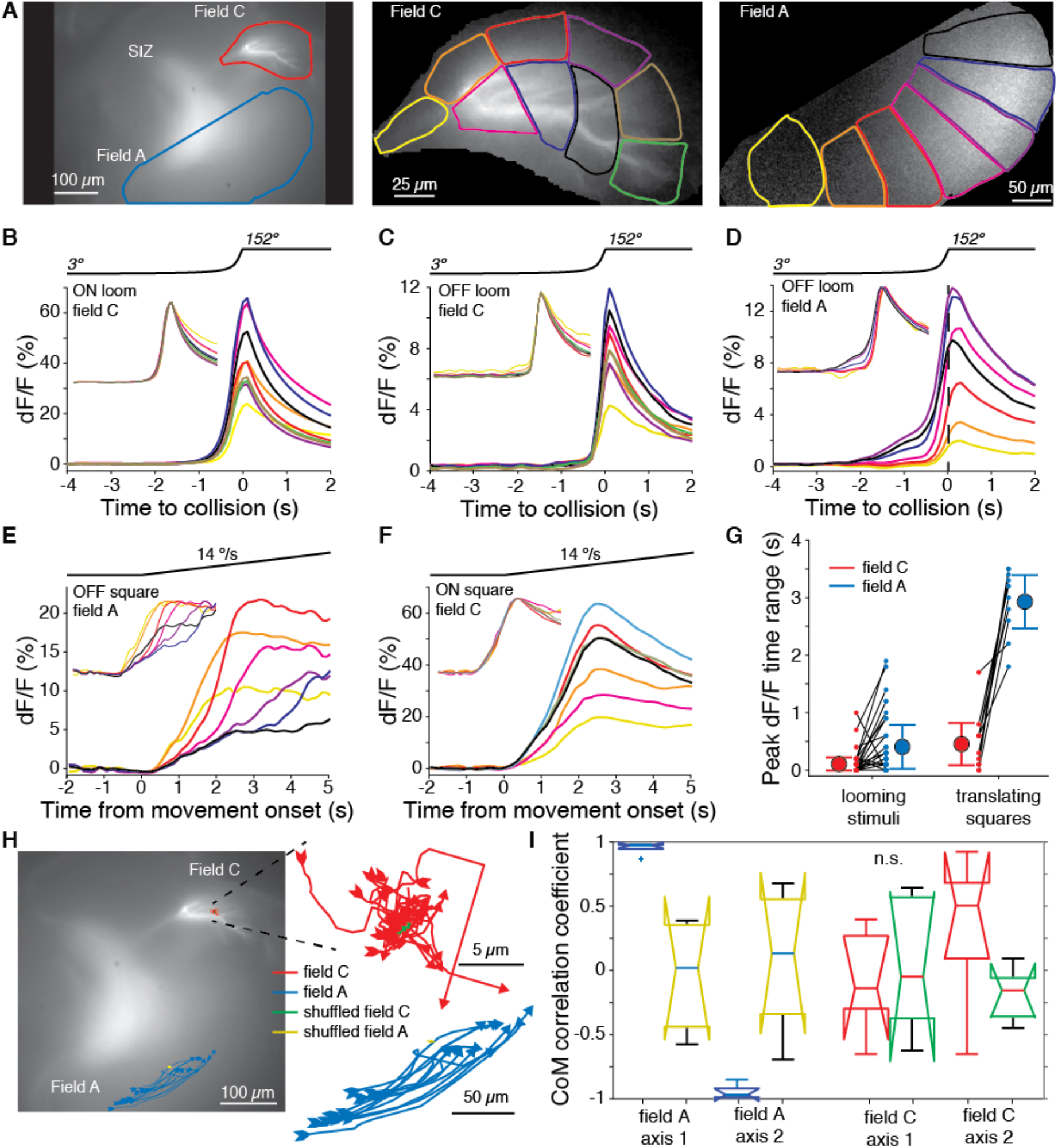
Within field comparisons show lack of retinotopy in field C. A) *Left:* Example dF micrograph in response to a ON loom with dendritic fields marked. *Middle:* Close-up of field C indicating the color-coded subregions used for dF/F calculations. *Right:* Similar close-up of field A in response to a black loom. B) Time course of mean dF/F in field C subregions shown in A in response to ON looms. The traces are rescaled to have the same peak value in the inset. C) Black looms elicited similarly timed responses across field C subregions. D) Black looms produced earlier and larger responses in field A subregions receiving inputs from the center of the loom. E) Responses to black translating squares show sequential activation of field A subregions (inset, as in B). F) ON translating squares produced a large synchronous dF/F signal across field C (inset, as in B). G) The range in peak dF/F loom response times between subregions was higher in field A than C (p = 3.2 • 10^−4^, t-test, N = 21). For translating squares, the range of dF/F peak times was larger for field A (p = 0.002, t-test, N = 8). H) Micrograph shown in (A) with superimposed trajectories of the dF/F center of mass in dendritic fields A and C for each trial of a translating bar (blue and red lines). For each dendritic field the center of mass trajectories of randomized data shown with yellow and green lines. At right, the same trajectories are shown zoomed in. I) The correlations between dF/F center of mass and bar position of each animal. Data from fields A and C are shown in blue and red respectively, and randomized data for each is shown in yellow and green. For H and I, data is taken from 21 trials of 5 animals.

To further test whether the field C mapping was random, we computed the correlation between the response center of mass (CoM) and stimulus position for small translating stimuli. For field A, the CoM of dF/F moved across the dendritic arbor as the stimulus traversed the visual field (Fig. 4H, blue lines). For all trials tested, the CoM position was correlated to stimulus position (p < 0.001). For field C, 9 of 21 trials had significant correlations between stimulus position and response CoM, but the CoM trajectories were short and inconsistent across trials within the same animal (Fig. 4H red lines). We compared these CoM trajectories to those of pixelwise randomization of the response (Fig. 4H, green and yellow lines). The field C CoM trajectories were not significantly more correlated with stimulus position than their randomly shuffled versions (Fig. 4I). The axis within field C that showed the strongest influence of stimulus position only had a mean shift in CoM of 3.5 µm (s.d. = 5.6) with a 115° change in stimulus position (Fig. 4H, right). These data indicate that there is no retinotopic mapping of field C excitation and the input locations are not different from random.

### Absence of ON spatial coherence sensitivity in LGMD and behavior

What are the consequences of the random mapping of ON synaptic inputs to the LGMD? Earlier work showed that both the retinotopic mapping of excitatory inputs and active dendritic processing in field A are critical for grasshoppers’ ability to discriminate the spatial coherence of black looming stimuli, a computation akin to object segmentation (Dewell and Gabbiani, 2018a). As the retinotopy and active conductances of field A are absent in field C, we used electrophysiology to test whether this prevented discrimination of the spatial coherence of white looms. For this purpose, we used the same approach as previously used for black stimuli (Dewell and Gabbiani, 2018a). We first pixelated the screen at the spatial resolution of photoreceptor receptive fields and replaced local edge motion by an equivalent local luminance change in each pixel to obtain ‘coarse looming stimuli’ (Fig. 5A, middle). We then randomly displaced each pixel at increasingly distant locations to obtain stimuli of decreasing coherence (Fig. 5A, bottom). Indeed, and in contrast to black looms, the peak firing rate of the LGMD did not change with the spatial coherence of white looms. However, as observed for black looms the responses contained more bursts for incoherent looms (Fig. 5B; Dewell and Gabbiani, 2018).

**Figure 5.**
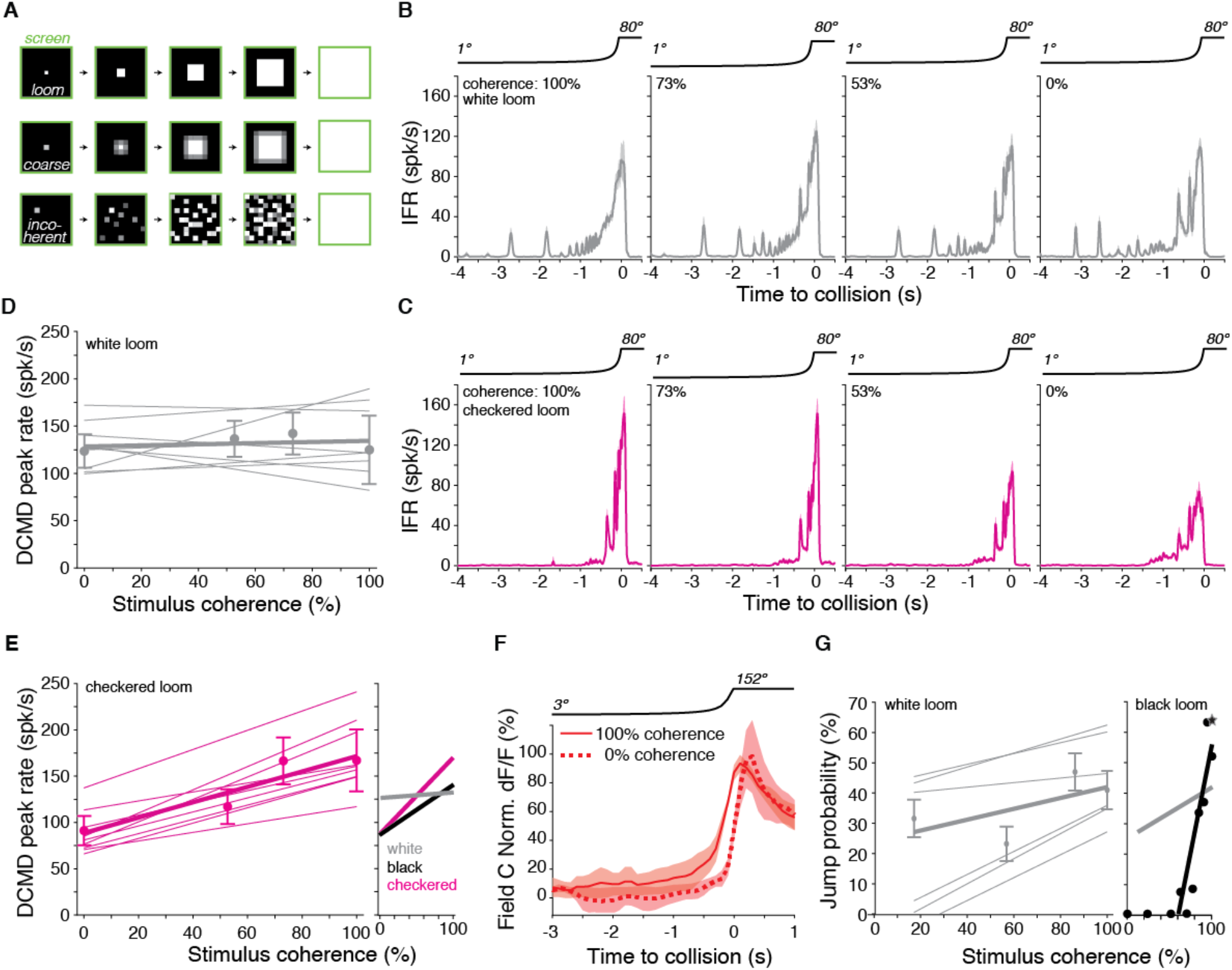
Lack of spatial coherence preference for white stimuli. A) *Top:* Schematic of looming stimulus. *Middle:* coarse looming stimulus. *Bottom:* stimulus with reduced spatial coherence. B, C) Firing frequency of the LGMD in response to white and checkerboard looming stimuli, respectively. Lines and shaded region are mean ± s.e.m., N = 9. As the spatial coherence was decreased, the response to checkered looms decreased (magenta). Responses to white looming stimuli did not decrease with reduction in spatial coherence (grey). D, E) Linear fits of the peak firing rate to the spatial coherence of the stimulus. Thin lines are fits to individual animals, thick lines are fits to the population, and points and error bars are population mean ± mad. Checkered stimuli had a mean ρ of 0.91, while for white stimuli, mean ρ was 0.15. Right inset: Firing rate as a function of stimulus coherence for black looms (from Dewell and Gabbiani, 2018) compared to data for white and checkered looms at left and in (D). F) Time course of field C normalized dF/F in response to 100% and 0% coherent white looming stimuli shows no change in peak value (N = 6). G) Left: Jump probability as a function of stimulus coherence for white looms (mean ± s.d., N = 6). Thick grey line: linear fit; thin grey lines: fits on individual animals. Right inset: Jump probability as a function of stimulus coherence for black looms (from Dewell and Gabbiani, 2018). Thick grey line is reproduced from left plot for comparison.

This difference in stimulus coherence sensitivity for white and black looms raised the question of how the LGMD would respond to stimuli containing a mixture of ON and OFF polarities, such as checkerboard stimuli (Fig. 1A). For spatially coherent stimuli, the peak firing rate for checkered stimuli was higher and occurred later than for solid white stimuli (p = 0.001, sign rank test). Reducing the coherence of black and white checkered looming stimuli decreased LGMD firing (Fig. 5C), as observed for solid black looms. For the nine animals studied only three responded maximally to the 100% coherent stimulus for white looms (Fig. 5D), while all nine animals responded maximally to the 100% coherent stimulus for checkered looms (Fig. 5E). Thus, the spatial coherence preference for checkered looms was similar to that of black looms characterized by Dewell and Gabbiani (2018a; Fig. 5E, right inset). The peak calcium fluorescence in field C was also unchanged by the spatial coherence of ON looms (Fig. 5F).

In response to black looms, the jump escape probability of grasshoppers is exquisitely sensitive to stimulus coherence and tightly correlated with the peak firing rate response of the LGMD (Dewell and Gabbiani, 2018a). To determine if the grasshoppers’ escape behavior depended on the spatial coherence of white stimuli, we presented the same stimuli to freely moving animals. The jump probability showed a slight decrease with reduced spatial coherence, but nothing like the steep coherence preference seen previously for black looming stimuli (Fig. 5G; Video 1 and 8). Thus, this reduced ability to discriminate the spatial coherence of an approaching white stimulus is presumably due to the excitatory ON inputs impinging non-retinotopically onto field C and a lack of active conductances devoted to processing them there (Fig. 4).

### Similar LGMD calcium responses for black and various mixed ON/OFF stimuli

So far, we have seen that ON stimuli excite dendritic field C non-retinotopically (Fig. 2), which leads to a lack of selectivity for the spatial coherence of ON looms (Fig. 5). But many real-world predators would contain a mix of light and dark regions, which would excite both dendritic fields. To measure how stimuli containing both ON and OFF inputs are processed, we imaged the calcium influx produced by approaching dark and light checkerboards and concentric squares on 50% luminance backgrounds (Fig. 6A, B). The dF/F was higher in field A than in field C during approach for both stimuli and the post-loom flash response was higher in field C (Fig. 6C, D), similar to the responses for solid black squares on a white background (Fig. 2F). For the seven animals tested, there was no difference in peak response between concentric squares, checkerboards, and OFF stimuli in either dendritic field (Fig. 6E, F). As in the earlier data, ON stimuli produced a higher peak response in field C and a lower peak response in field A (p = 0.015). The response at the SIZ showed no preference for ON, OFF, or mixed stimuli indicating similar spiking output from all four tested stimuli (Fig. 6G).

**Figure 6.**
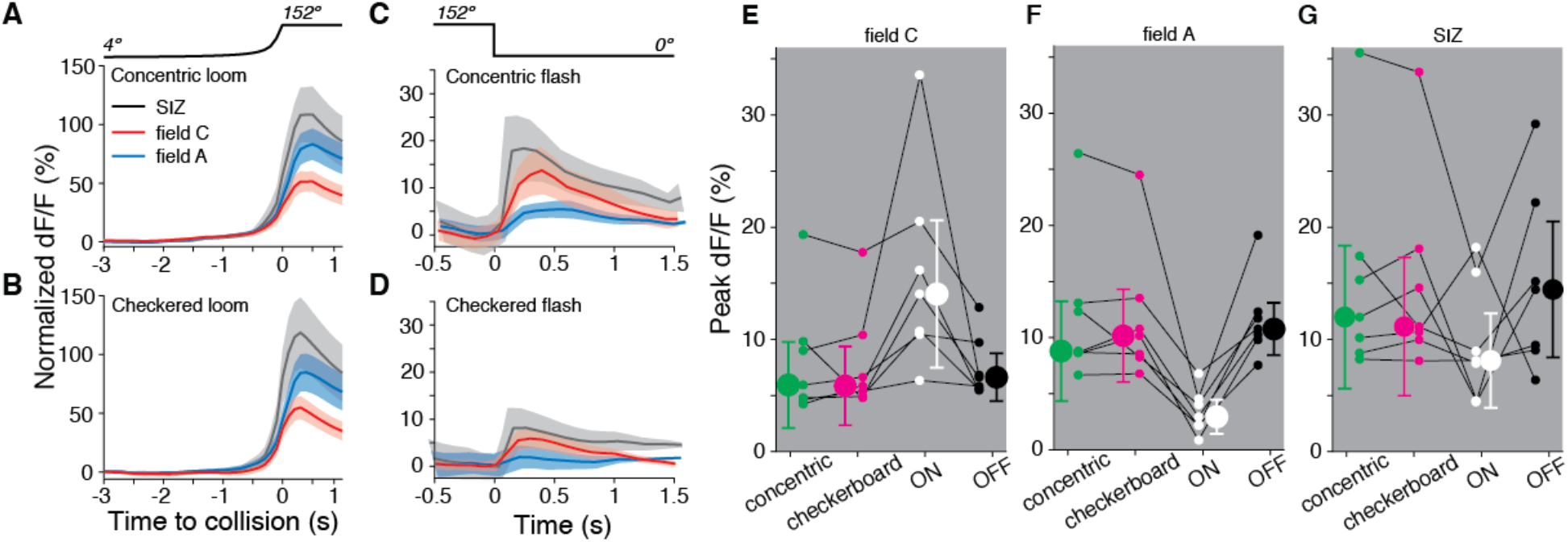
Both dendritic fields A and C respond similarly to looming stimuli of mixed luminance and to black looms. A) Looming stimuli composed of 5 concentric ON and OFF squares produce slightly higher responses in field A (p = 0.016, ST). B) Checkered looming stimuli similarly produced higher field A responses (p = 0.016, ST). C, D) Post loom flash responses were higher in field C after concentric looming stimuli (p = 0.026, ST) but not for post-checkered flashes (p = 0.093, ST). E) Field C peak dF/F for each stimulus. ON looms elicited the highest response in all animals (p = 0.015, ST), and responses to the other 3 stimuli were not significantly different (p = 0.82, KW). F) Field A peak dF/F for each stimulus. ON looms elicited the lowest response in all animals (p = 0.015, ST), and responses to the other 3 stimuli were not different (p = 0.89, KW). G) All stimuli produced similar peak dF/F at the SIZ (p = 0.26, KW). In A-G, N = 7 animals. ST: sign test; KW: Kruskal-Wallis test.

### Energetic implications of ON/OFF dendritic mappings

To further explore the functional significance of segregating ON and OFF excitation between dendritic fields, we conducted a series of simulations on a biophysical model of the LGMD neuron. Previous LGMD models have reproduced many properties of the neuron, including responses to black looming stimuli (Dewell and Gabbiani, 2018a). The current model used the same pattern of synaptic inputs and model morphology (Fig. 7A), with updated membrane parameters incorporating findings from subsequent investigations (Dewell and Gabbiani, 2018b, 2019).

**Figure 7.**
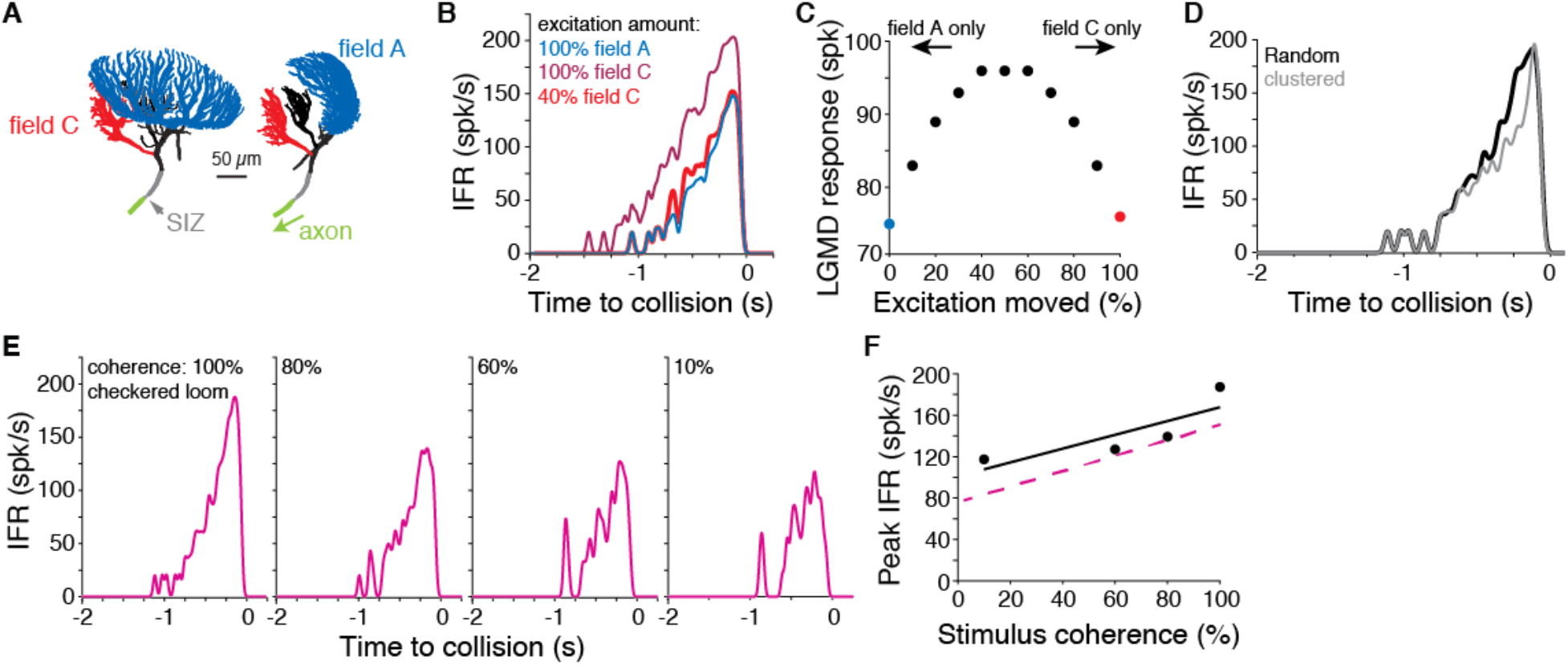
Simulations of a biophysical LGMD model reveal energetic savings of segregated inputs. A) Images of the model morphology showing the dendritic fields; the right image shows the model neuron rotated 90° from the left image. B) Instantaneous firing rate (IFR) of the model’s response to a black loom with excitation impinging on field A (blue line; same synapse locations as in Dewell & Gabbiani 2018). Moving all excitation to random field C locations (dark red) increased firing. Removal of 60% of excitatory synapses still produced a response as high as the field A inputs (red). C) If excitatory synapses of simulated looms were distributed between the dendritic fields (with removal of 60% of synapses moved to field C), responses were highest with inputs split evenly between fields. Red and blue points are same as simulations shown in (B). D) If all excitation impinged on field C with excitatory inputs clustered and spreading from a central point to replicate a retinotopic mapping, responses were reduced from inputs with random mapping. E) Simulations reproduced firing patterns of response to checkered looming stimuli of different spatial coherences (c.f. Fig. 5C). F) Spatial coherence preference for checkered looms for the model (black) and experimental data (dashed magenta).

The first simulation tested how inputs to fields A and C differ in their ability to initiate action potentials. Excitation to field A generated by a simulated black loom produced reliable firing (Fig. 7A, B, blue). Moving the excitatory inputs from field A to C produced much higher firing (Fig. 7B, dark red). This was due largely to the relative electrotonic proximity of field C to the SIZ; consequently, the depolarization elicited by synaptic inputs in field C attenuates less as it propagates towards the SIZ. Reducing the number of field C excitatory synapses by 60% without changing the amount of inhibition still produced as much firing as the field A excitation simulation.

Calcium fluorescence showed that looms of all contrasts produced some excitation in both dendritic fields (Fig. 2), so we next tested how the LGMD response changed with excitation split between the fields. For simulated looming stimuli, we varied the fraction of excitation distributed between the dendritic fields while removing 60% of the field C inputs since this achieved the same firing output regardless of dendritic field input assignment (see Fig. 7B). The more evenly the inputs were split between the fields, the larger the resulting neural response was (Fig. 7C). Thus, an even splitting of excitation between the fields A and C maximizes its effectiveness at triggering LGMD spiking.

ON synaptic inputs show no retinotopy and are distributed randomly throughout dendritic field C (Fig. 4). Yet, dendritic field C is sufficiently large electrotonically to allow for a retinotopic mapping similar to that observed in field A. To test if the lack of field C retinotopy changed the LGMD’s output we simulated clustered excitatory inputs that spread across field C as would occur with a retinotopic mapping. We found that such a mapping led to reduced firing to looming stimuli (Fig. 7D). Similar results were observed in field A when the active membrane conductances responsible for coherence discrimination were removed (Dewell and Gabbiani, 2018a). Thus, in the absence of additional processing by active membrane conductances to implement coherence selectivity, random synaptic localization of ON inputs in field C is most effective at triggering LGMD spiking. This arrangement minimizes the reduction in synaptic current resulting from a decrease in driving force when adjacent locations in visual space are stimulated and mapped to nearby dendritic locations.

The LGMD remains sensitive to the spatial coherence of checkered stimuli despite a substantial fraction of excitation impinging on field C (Fig. 5). To test whether the model reproduced this result, we simulated checkered looms of different coherence levels. In these simulations excitation from OFF checkered regions impinged on field A and ON regions excited field C. The model reproduced the decreased firing and increased bursting of spatially incoherent stimuli seen in experimental data (Fig. 7E). The overall spatial coherence preference was qualitatively similar in model and experiment (Fig. 7F). This confirms that for these mixed contrast stimuli, the retinotopic mapping and active filtering of field A can produce spatial coherence selectivity even when 50% of the excitatory inputs are moved out of field A. Taking into account the 60% reduction in the required excitatory inputs to field C due to their proximity to the SIZ, this represents a 30% total input reduction without a loss in sensitivity to spatial coherence for textured stimuli. As solid black looming stimuli partially excite field C (Fig. 2), these simulations suggest energetic savings of ∼10% for black stimuli while generating the same spiking output. Stimuli with a mix of ON and OFF inputs would also benefit from higher savings than if all excitation impinged on field A.

## Discussion

### Dendritic segregation of ON/OFF excitation shapes looming detection and escape behavior

We investigated a looming-sensitive neuron that integrates ON and OFF inputs and responds to approaching objects with a characteristic firing profile invariant to object contrast (Simmons and Rind, 1997; Gabbiani et al., 2001). Although, the LGMD neuron responds to either white or black approaching objects, whether grasshoppers escape to both was previously untested. We found that animals escaped from simulated white objects approaching on a collision course (Fig. 1) even if their spatial coherence was removed (Fig. 5). The behavioral and physiological responses generally occurred earlier for ON than OFF or checkered stimuli (Figs. 1, 2). The ability to detect and respond to white objects but not to discriminate their spatial coherence is due to a previously unknown segregation of ON and OFF excitation and their distinct mapping onto separate dendritic fields.

Unlike the previously described OFF excitatory inputs which are retinotopically mapped with high precision on dendritic field A, the ON excitatory inputs impinge non-retinotopically onto field C (Fig. 4). Both the retinotopic input mapping and the active conductances within field A are critical for OFF spatial coherence selectivity (Dewell and Gabbiani, 2018a; Zhu et al., 2018). Although it is not known whether field C is completely passive, it lacks the HCN channels present in field A which were found to be necessary for the animal’s behavioral selectivity to the spatial coherence of black approaching stimuli (Dewell and Gabbiani, 2018a). Thus, our results demonstrate the computational and behavioral consequence of distinct dendritic synaptic input mappings and active conductance localization.

As real approaching predators likely exhibit non-uniform visual contrast, we also tested responses to stimuli with a mix of ON and OFF regions. Checkered stimuli produced responses very similar to solid black stimuli (Figs. 5, 6). The LGMD was responsive to checkered stimuli and discriminated their spatial coherence, demonstrating that only part of the excitation needs to impinge retinotopically onto field A to evoke selective escape responses. Visual detection of approaching predators is believed to contain more OFF than ON information (Zhou et al., 2022). so this neural and behavioral selectivity for the spatial coherence of black and checkered stimuli but not white ones likely indicates that grasshoppers can discriminate the spatial patterns of real-world threats.

Although we cannot experimentally test the behavioral response of an animal lacking field C from the LGMD, another looming-sensitive neuron laying adjacent to the LGMD called the LGMD2 provides some insight. The LGMD2 has a dendritic field A of the same size and shape as that of the LGMD but lacks analogous fields B or C. It responds with a strong preference to dark objects and exhibits limited ability to detect white approaching stimuli (Simmons and Rind, 1997; Rind and Leitinger, 2000). This further suggests the necessity of the additional dendritic field and its ON excitatory inputs for the LGMD’s detection of impending collisions independent of contrast polarity.

### Interpretation of calcium fluorescence

Both ON and OFF pathways excite the LGMD through calcium-permeable nAChRs (Fig. 3; Peron et al., 2009). This allowed characterization of the functional segregation between fields with an intracellularly-injected fluorescent calcium indicator (Methods). The exact relationship between the amount of synaptic excitation and dF/F remains unknown but does not affect our interpretation of experimental results. The intracellular injection of the fluorescent dye typically yields an uneven fluorescence distribution within and between cells. Thus, the dF/F values indicate a general activity level of the dendrites, dependent on a combination of pre- and post-synaptic properties. For black looming stimuli, the dF/F is well correlated to the subthreshold membrane potential (Zhu et al., 2018).

### Visual processing of ON and OFF pathways

In mammals and insects, visual processing is similarly split between ON and OFF pathways. In both cases, photoreceptors respond to luminance changes of either polarity but at the next stage increments and decrements are encoded by different neurons – ON and OFF bipolar cells in the mammalian retina and L1 and L2 cells in the lamina of the insect optic lobe (Clark and Demb, 2016). The rectification of L1 and L2 neurons is not complete, though, causing the ON and OFF pathways to respond weakly to luminance changes of the opposing polarity (Clark et al., 2011; Strother et al., 2014). Hence, the most parsimonious explanation for responses of dendritic fields A and C to white and black stimuli is that they receive only OFF and ON inputs, respectively, but that neither pathway is wholly selective to contrast polarity.

In both insects and mammals the OFF pathway is more sensitive to fast movement than the ON pathway (Leonhardt et al., 2016; Mazade et al., 2019). In mammals OFF neurons are biased towards central vision (Mulholland and Smith, 2021; Williams et al., 2021). In insects, the columnar organization of the optic lobe likely produces an equal distribution of ON and OFF neurons. The central bias in mammals is related to the smaller receptive fields of OFF neurons (Mazade et al., 2019). In the grasshopper, the excitatory OFF inputs to field A of the LGMD have the same spatial resolution as the eye (∼2º; Jones and Gabbiani, 2010). Feedforward inhibitory inputs to field C have lower resolution with receptive fields 10-fold larger (Wang et al., 2018; Zhu et al., 2018). The similarities of contrast polarity encoding between taxa likely result from visual processing being tuned to the statistics of natural scenes (Clark et al., 2014; Clark and Demb, 2016; Chen et al., 2019).

As this is the first report of ON excitation to field C, the inputs have not yet been characterized and little is known about the pre-synaptic circuitry. The known anatomy and current data suggest much fewer presynaptic neurons with larger receptive fields than in field A (Elphick et al., 1996; Zhu et al., 2018). The dorsal uncrossed bundle (DUB) that projects to field C contains ∼500 neurons and was previously believed to only convey feedforward inhibitory input. Since inhibitory input is conveyed by a small subset of these neurons (Wang et al 2018), most DUB neurons are likely conveying the ON excitation described herein.

### Energy costs of looming detection

The nervous system is an energetically expensive organ, estimated to consume over 20% of calories in humans. Reducing the cost of neural processing is thus critical for retinal and cortical circuits (Vincent and Baddeley, 2003; Hasenstaub et al., 2010). Excitatory synaptic transmission contributes greatly to this energetic cost, likely accounting for more than 40% of the brain’s total ATP consumption (Sibson et al., 1998; Attwell and Laughlin, 2001). Most of that expense comes from the pumping of Na^+^ and Ca^2+^ ions out of neurons following excitatory synaptic events (Attwell and Laughlin, 2001; Hasenstaub et al., 2010). The LGMD has over 100,000 excitatory synapses which pass Na^+^ and Ca^2+^ (Peron et al., 2009; Rind et al., 2016). Thus, reducing the number of synaptic events needed to detect an approaching object could potentially generate large energetic savings. Indeed, based on estimates of ATP cost per excitatory synaptic event (Attwell and Laughlin, 2001), our estimates for the number of excitatory synaptic events per loom, and the estimated saving of 60% excitation with a move from field A to C suggested by our simulations (Fig. 7) ∼10 billion ATP molecules could be saved per looming response. The LGMD receives inputs from the full field of view with a high rate of visually-elicited and spontaneous excitation, so the energetic savings would not be limited to periods of synaptic excitation caused by approaching predators. This estimate assumes that the pre-synaptic circuitry for ON and OFF inputs are equally energetic and neglects any energy expense differences in dendritic processing. As the ON inputs likely comprise fewer neurons and field A has more active conductances, energetic savings are likely higher than estimated here. These energetic savings are due primarily to inputs impinging closer to the site of spike initiation.

### Functional significance of synaptic mapping within dendrites

The additional energetic costs of OFF excitation to field A raises the question of why the LGMD wouldn’t have all inputs excite the more proximal field C. The proposed answer lies in the stark difference in selectivity of responses to ON and OFF stimuli. While the LGMD responds to white looms and the animals jump to escape them, the response is not selective to the spatial coherence of the stimulus (Fig. 5). As escaping predation requires not just detecting threats, but discriminating them from non-threatening stimuli, spatial coherence selectivity is likely critical for survival. As of yet, there has been no examination of whether mammalian looming-detection circuits are sensitive to stimulus spatial coherence or how they integrate inputs from ON and OFF pathways. Mice have a stronger behavioral response to black looms and looming-sensitive neurons in the superior colliculus show a preference for OFF inputs (Yilmaz and Meister, 2013; Branco and Redgrave, 2020).

Within the LGMD the OFF pathway excites in precise retinotopic manner a distal dendritic field with large HCN and inactivating K^+^ conductances that enable discrimination of spatiotemporal input patterns (Zhu and Gabbiani, 2016; Dewell and Gabbiani, 2018a). Conversely, the ON pathway results in non-retinotopic excitation of a proximal dendritic field lacking these active channels. Voltage-gated Ca^2+^ channels, Ca^2+^-dependent K^+^ channels, and M-type K^+^ channels that control bursting and spike-frequency adaptation are located close to the SIZ (Peron and Gabbiani, 2009; Dewell and Gabbiani, 2018b). The more distal position of field A makes integration within its dendrites further removed from the influence of these conductances and more dependent on the channels localized within it. The increased electrotonic distance from the SIZ also increases the resolving power of dendritic integration in response to synaptic input patterns, since more excitatory inputs are required to generate action potentials.

The combination of reduced energy cost of field C inputs with increased discrimination by active processing in field A suggests a possible general principle for dendritic mapping whereby finer discriminations of the synaptic pattern are implemented in distal, active dendrites while coarser discriminations are implemented by proximal dendrites. Additionally, having excitatory inputs spread between dendritic regions offers energetic savings of its own as concentrating inputs reduces the driving force of the activated receptor channels (Fig. 7C, D). Examination of integration in neocortical or hippocampal pyramidal neurons which contain distinct dendritic regions and segregated inputs may provide a test for the generality of this mapping principle.

Within hippocampal pyramidal neurons, distal CA1 dendrites receive excitation from entorhinal cortex while proximal dendrites are excited by inputs from CA3. These dendritic regions also have distinct sets of active channels (London and Hausser, 2005; Spruston, 2008; Lefebvre et al., 2015). An alternative hypothesis to that put forth above suggests that proximal excitation is the primary driver of spiking while distal excitation is modulatory (Behabadi et al., 2012; Hawkins and Ahmad, 2016). The two ideas are not incompatible, though. Active processing in distal dendrites of cortical pyramidal neurons can produce fine discrimination of the spatiotemporal pattern of excitatory inputs, even if dendritic compartmentalization and the location-dependence of synaptic integration are variable (Poirazi and Papoutsi, 2020).

Within cortex, feedback from higher cortical areas excite the distal apical dendrites of pyramidal neurons which also have increased active conductances that can produce fine discrimination of distal inputs (Schuman et al., 2021). It has been suggested that the dendritic segregation of feedforward and feedback cortical inputs might underlie the comparison of sensory (bottom-up) inputs with top-down predictions (Larkum, 2013). In this context, our work suggests that distal feedback inputs might allow a finer discrimination of top-down than proximal bottom-up inputs. Within each neuron type, evolution presumably constrains the energetic costs of neural signaling required for dynamic, nonlinear computations. Systems in which we can simultaneously examine membrane properties, behavioral function, and dendritic mapping are crucial for discovering the underlying biophysical mechanisms.

## Methods

### Preparation

The experimental procedures used have been previously described (Dewell and Gabbiani, 2018a; Zhu et al., 2018). Experiments were conducted on adult *Schistocerca americana* grasshoppers 1 to 4 weeks after their final molt. Preference was given to larger females ∼3 weeks after final molt that were alert and responsive. Animals were selected for health and size without randomization. For calcium imaging, the head was opened to expose the brain and optic lobes. After removing the sheath protecting the right optic lobe, the calcium indicator Oregon Green Bapta (OGB-1) was injected by iontophoresis into the LGMD with sharp intracellular electrodes (Zhu and Gabbiani, 2016; Zhu et al., 2018). The amplitude and duration of current steps for OGB iontophoresis were manually adjusted to produce a dim stain of the entire dendritic arbor. The variability in injection locations and baseline fluorescence levels produced a wide range of signal strengths across animals, with lower baseline fluorescence producing larger signal-to-noise ratios under visual stimulation.

### Imaging

Calcium fluorescence was imaged with a CCD camera as previously described (Zhu et al., 2018). Images were captured through with a 16X/0.8 numerical aperture (NA) water immersion objective lens for imaging (CFI75 LWD 16XW, Nikon Instruments) and saved at 5 Hz with a Rolera XR camera (Qimaging, Surrey, BC, Canada). The image resolution was 696×520 and saved as lossless 12-bit motion jpeg movies. The spatial resolution of the images was 0.9 µm per pixel.

### Visual stimulation

For calcium imaging experiments, visual stimuli were generated with the Psychtoolbox (PTB-3) and MATLAB (MathWorks, Natick, MA) as done previously (Zhu et al 2018). A digital light processing projector (DLP LightCrafter 4500, Texas Instruments, Dallas, TX) displayed stimuli on a screen (non-adhesive stencil film, 0.1 mm thick) placed 20 mm from the right eye. The right eye was exposed, and the head and neck were submerged in saline. A mechanical brain holder was placed under the optic lobe to prevent movement perpendicular to the imaging plane. The DLP was programmed in pattern sequence mode to display 6-bit green-scale images with a refresh rate of 240 Hz and 912×1140 resolution. For behavior and electrophysiology data shown in figures 1 and 5, visual stimuli were generated by custom C software from a QNX4 computer and displayed on a CRT monitor with a 200 HZ refresh rate at 640×480 resolution (Gabbiani et al., 1999; Dewell and Gabbiani, 2018a). Stimuli for all experiments were displayed with 6-bit resolution luminance values. For behavioral experiments looming stimuli had a *l*/|*v*| of 40, 80, or 120 ms, and for calcium imaging stimuli with *l*/|*v*| = 100 or 120 ms were used. Spatial coherence of looming stimuli was reduced by pixelating the stimuli into 2.5° regions and adding a spatial jitter to these coarse pixels (Dewell and Gabbiani, 2018a). For behavior test of coherence selectivity the stimulus *l*/|*v*| = 80, and for the DCMD recordings *l*/|*v*| = 50.

### Behavior experiments

Jump experiments were conducted as previously (Fotowat and Gabbiani, 2007; Dewell and Gabbiani, 2018a). Videos were recorded with a high-speed digital video camera (GZL-CL-22C5M; Teledyne Flir), equipped with a variable zoom lens (M6Z 1212–3S; Computar, Cary, NC). Image frames were recorded at 200 frames per second with the acquisition of each frame synchronized to the vertical refresh of the visual stimulation display (Xtium-CL PX4; Teledyne Flir). Videos were made from the 12-bit images and saved in lossless motion JPEG format using custom Matlab code.

### Data analysis and statistics

All data analysis was carried out with custom MATLAB code (MathWorks, Natick, MA). For calcium fluorescence, data was saved in either uncompressed AVI or lossless motion JPEG format using Matlab’s Image Acquisition toolbox. Most trials had no motion artifacts, but when present, translational motion artifacts were corrected by aligning dendritic locations across frames using the “imregcorr” function of Matlab’s Image Processing toolbox. After any motion correction, videos were median filtered with a 3×3 pixel window to reduce noise. Fluorescence change (dF) was normalized to the baseline fluorescence, calculated as the average value in the 2 seconds before stimulus presentation to calculate dF/F. In some trials, the baseline fluorescence changed over time even in the absence of visual stimuli and a linear fit to the fluorescence time course before stimulus presentation was subtracted instead of a single value. Every trial was manually checked to confirm the change in fluorescence before any further data processing.

All regions of interest (ROIs) were drawn freehand and the reported dF/F is the mean value of all pixels within the selected region. The period from the start of looming stimuli until 1 s after the end of expansion was considered the looming response. For post-loom flashes, the peak dF/F of the flash response was measured as the difference between the maximum dF/F within 1.5 seconds after the flash minus the minimum dF/F in the 1 second before the flash.

For testing whether there was a retinotopic mapping of inputs in field C, the center of mass of the dF/F within a dendritic field was calculated for each frame while the bar was moving using Matlab’s image processing toolbox (using the “weighted centroid” property of “regionprops”). The trajectory of the center of mass was then compared to the stimulus position at each corresponding time. To test the randomness of this mapping, each pixel of the dF/F within the dendritic field was randomly redistributed and the center of mass trajectory was recalculated.

For statistical tests between jump probabilities, we used Fisher’s test to compare responses across stimulus speed and contrast. The timing of jump behavior was tested with the Wilcoxon rank-sum test. Paired comparisons of peak dF/F between dendritic fields or within dendritic fields for different stimulus polarities were made using a two-sided sign test. Comparisons across more than two stimuli (Fig. 6) were made with the Kruskal-Wallis test, a non-parametric version of the classical analysis of variance (indicated by KW in text). All correlations were computed using Pearson’s linear correlation coefficient.

## Acknowledgements

We would like to thank Drs. Alyse Thomas and Jake Reimer for manuscript feedback. This work was supported by grants from the National Science Foundation (DMS-1120952 and DBI-2021795) and NEI Core Grant for Vision Research (EY-002520-37).

## Supplemental movies

Video 1. Example jump escape response of a grasshopper to a white looming stimulus. As the stimulus expands, the animal jumps and flies away before the time of projected collision. Frame rate: 200 Hz; slowed to 30 Hz. Compare to a similar response for a black looming stimulus in Video 3 of Dewell & Gabbiani (2018a); DOI: 10.7554/eLife.34238.008.

Video 2. Example of raw fluorescence signal within the LGMD in response to a white looming stimulus. As the looming stimulus expands, note the fluorescence increase in field C (top right, in focus). Baseline field A fluorescence is also visible at the center. Frame rate for videos 2-7: 5 Hz; real speed. Matches data of Fig. 2C,E. Image size for videos 2-7: 630 × 470 µm.

Video 3. The calculated fluorescence dF/F of the same trial shown in Video 2. Note the disappearance of field A baseline fluorescence through this manipulation.

Video 4. An example of the fluorescence dF/F during a black looming stimulus. Data is from the same LGMD as shown in videos 2 and 3. Matches data of Fig. 2B,D.

Video 5. Example of raw fluorescence signal within the LGMD in response to a white moving dot stimulus. As the object moves, the fluorescence increases in all of field C (top center, in focus). Compare to similar response to a black dot in Video S2 of Zhu *et al*. 2018; https://www.cell.com/cms/10.1016/j.celrep.2018.04.079/attachment/74ca6771-85e8-43e1-8804-9367a439e612/mmc3.mp4

Video 6. The calculated fluorescence dF/F of the same trial shown in Video 5. Matches data in Fig. 4F.

Video 7. An example of the fluorescence dF/F during a black moving dot stimulus. Data is from the same LGMD as shown in videos 5 and 6. Note strong activation in field A sweeping along a crescent starting from bottom center (out of focus). In addition, some activation is also seen in field C (top right, in focus). Matches data in Fig. 4E.

Video 8. Example response showing a grasshopper jump escape in response to a white 20% spatially coherent looming stimulus.

